# Structural and functional reference maps of transcriptional enhancers in the developing human cortex

**DOI:** 10.64898/2026.11.24.768903

**Authors:** Druha Karunakaran, Kelsey Hennick, Rachel Leonard, Tomasz Nowakowski, Hanna E Berk-Rauch, Aravinda Chakravarti

## Abstract

Non-coding regulatory variation is the major driver of human phenotypic diversity, yet maps of cis-regulatory elements (CREs) and their variability across developmental cell types remain limited. Here, we integrated single-nucleus chromatin accessibility data from 54 mid-gestation human prenatal cortical donors to generate a CRE atlas comprising 1,165,697 candidate CREs across 104,266 cortical nuclei. Chromatin-state annotation classified 1.5% of CREs as bivalent promoters, 0.3% as active promoters, 39.8% as active distal enhancers, 1.3% as CTCF-associated insulators but 57.1% remain unresolved. Overlap with an external enhancer assay dataset showed that many unresolved CREs, nevertheless, exhibit regulatory activity; thus, current chromatin signatures underestimate the functional regulatory landscape. Pseudo-bulk analysis across major cortical lineages revealed that the prenatal cortical regulome is dominated by lineage-restricted CREs, with only a small core shared across all lineages. Restricted CREs were enriched for unresolved elements, intermediate-sharing classes were enhancer-dominant, and broadly shared CREs showed the strongest promoter-proximal and bivalent contribution. Lineage-selective transcription factor (TF) sequence grammar analyses identified progenitor-associated homeodomain programs and neuronal basic helix–loop–helix signatures, together with representative candidate regulators such as SOX10, NFIA, EMX2, DLX1, ARX, and RFX3. Across donors, developmental time, and sex, prenatal cortical CREs segregated into a stable versus variable regulatory compartments. Broadly, recurrent and developmentally persistent CREs had greater activity, on average, and greater enrichment for promoter-associated states, whereas donor-restricted and temporally restricted CREs had lower activity and were enriched for unresolved and enhancer-like elements. Sex-biased CREs were rare but highly specific: female-biased CREs were predominantly X-linked, broadly shared, and developmentally persistent, whereas male-biased CREs were more numerous, less resolved, and more restricted across lineage and developmental contexts. Together, these results define the prenatal cortical regulatory landscape at single-cell resolution, revealing a small, stable, promoter-rich core and a larger variable, enhancer- and unresolved-rich compartment in which lineage diversification is driven primarily by selective deployment of distal regulatory elements.

## INTRODUCTION

Genome-wide association studies (GWAS) have convincingly demonstrated the ubiquity of non-coding sequence variation as the major causal driver of human phenotypic variation, including inter-individual disease risk.^1^ These studies have also uncovered the tremendous polygenicity of human phenotypic inheritance.^2^ The emergent genetic model is that sequence variation within mostly cis-regulatory elements (CREs), and sometimes their cognate transcription factors (TFs), which are far more conserved, alter the expression of target genes that are phenotypic contributors.^2–4^ Developmental^5^ and experimental^6^ data suggest that these causal genes are themselves co-regulated through one or more gene regulatory networks (GRNs), networks that display greater conservation in their functional architectures than their structures or sequences.^7^ These features can explain the specificity of both monogenic and polygenic phenotypes and how rare and common variants in target genes and their CREs affect phenotypic inheritance.^8^

Despite this compelling genomic model of complex trait inheritance, there are very few instances where multiple rare and common variants within a specific collection of genes in GRNs have been demonstrated to be *causal* for a phenotype, as well as explain *most of its variation*.^9–12^ The biggest impediment to such progress, except in rare circumstances, has been the availability of accurate genome wide maps of CREs, together with their target genes and GRNs, in specific cell-types and tissues. These are still in the early but active stages of development, despite the success of the NIH Roadmap^13^ and ENCODE^14,15^ projects. Even when available, such genome-wide maps are constructed from a limited set of samples so that the effects of genetic variation on this map cannot be assessed. The latter is crucial for phenotypic dissection because gene expression regulation is dynamic and sequence-dependent, and the basic premise of GWAS-based genetics is variation in CRE structure and function.^16,17^ A broad cross-tissue resource can tell us where candidate regulatory elements exist but understanding how they function together as networks within cells requires a single, developmentally coherent system. The fetal cortex is a particularly powerful tissue to test this hypothesis because it is highly dynamic, cell-type-rich, central to human developmental phenotypes, and strongly implicated in neurodevelopmental disease.

In this study, motivated by genomic analysis of autism spectrum disorder (ASD) and other neurodevelopmental disorders (NDDs),^18–22^ we investigate such variation in genome-wide CRE maps of the developing human fetal cortex. There is considerable evidence that both rare and common genetic variants in ASD are enriched in evolutionarily conserved CREs and, more specifically, within TF binding sites,^23–25^ and, that the mid-gestation cortex is a key contributor since ASD-associated genes show enriched expression in this tissue and developmental period, particularly in neuronal cell types.^18,26^ Beyond ASD and related NDDs, regulatory programs active during prenatal development of the cerebral cortex are likely relevant to a broader range of neuropsychiatric, cognitive, and neurological phenotypes, including schizophrenia, ADHD, epilepsy, and intellectual disability.^27–31^

In this study, we define cis-regulatory elements (CREs) operationally as accessible DNA elements identified by chromatin accessibility profiling^32,33^ and, where possible, annotated by chromatin-state information.^13,34,35^ Integration of accessibility with histone-marks and architectural-protein profiles enables many CREs to be classified as promoters, enhancers, or insulators, but these categories and classifications are incomplete. Promoter and enhancer functions can overlap, and some experimentally active regulatory elements lack canonical chromatin signatures altogether, such as instances of transcriptional ‘facilitators.’^36^ This limitation is particularly important in dynamic developmental tissues such as the cerebral cortex, where incomplete cell type- and stage-matched annotation may leave large portions of the functional regulatory landscape unresolved.

Here, we integrated epigenomic datasets from two studies^37–39^ to construct a comprehensive prenatal cortical CRE atlas^40^ and analyze it across multiple levels, including chromatin-state annotations, cell lineage sharing, transcription factor sequence logic, CRE– gene linkage, and variation across individuals, developmental time, and sex. Rather than simply expanding existing cell atlases, our goal was to define how regulatory elements are organized across developmental contexts and to identify which parts of the prenatal cortical regulome are stable versus variable. This framework provides a basis for interpreting non-coding regulatory variation in neurodevelopment and for generating hypotheses about mechanisms relevant to ASD and related neurodevelopmental disorders.

## RESULTS

### Comprehensive mapping of the developing cortical regulome

To determine the genomic regulatory architecture of the developing cerebral cortex, we analyzed newly generated single-nucleus RNA (snRNA-seq) and ATAC (snATAC-seq) Multiome profiles from 48 mid-gestation tissue donors,^39^ together with previously generated snATAC-seq data from 6 donors,^37^ comprising a total of 104,266 cells. Both datasets were generated from individuals around mid-gestation (GW16-24), a period coinciding with high levels of neurogenesis (Figure 1A, 1B).^38,41,42^ Because detailed regional annotations were not uniformly available across datasets, we treated all samples as ‘cortical’ and applied our published quality-control procedures to each dataset separately prior to integration.^43–47^ Using a unified indexing approach for chromatin accessibility reads,^40^ we harmonized peak coordinates across samples to generate non-redundant CRE maps for each dataset, termed the ‘Ziffra’^37^ and the ‘Hennick’^39^ maps, and a merged cortical reference map (Figure 1A). The Ziffra and Hennick maps contained 768,785 (median width 260bp based on 6 donors) and 702,326 (median width 304bp based on 48 donors) CREs, respectively, and the merged map comprised 1,165,697 CREs spanning 406 mb, or approximately 13% of the human genome. Chromosomally, the merged cortical map included 1,136,548 autosomal (97.5%), 26,651 X-linked (2.3%), 723 Y-linked (0.05%), and 1,775 pseudo-autosomal (0.16%) CREs. Of these, 128,244 (11%) overlapped coding sequence, 779,385 (67%) were genic but non-coding (defined as gene-associated and within 50kb of a gene boundary), and 258,068 (22%) were intergenic (Figure 1C). These results suggest that CREs are depleted on sex-chromosomes but enriched within exons and near genes. This non-random distribution is *a priori* evidence of their functional relevance.

**Figure 1.**
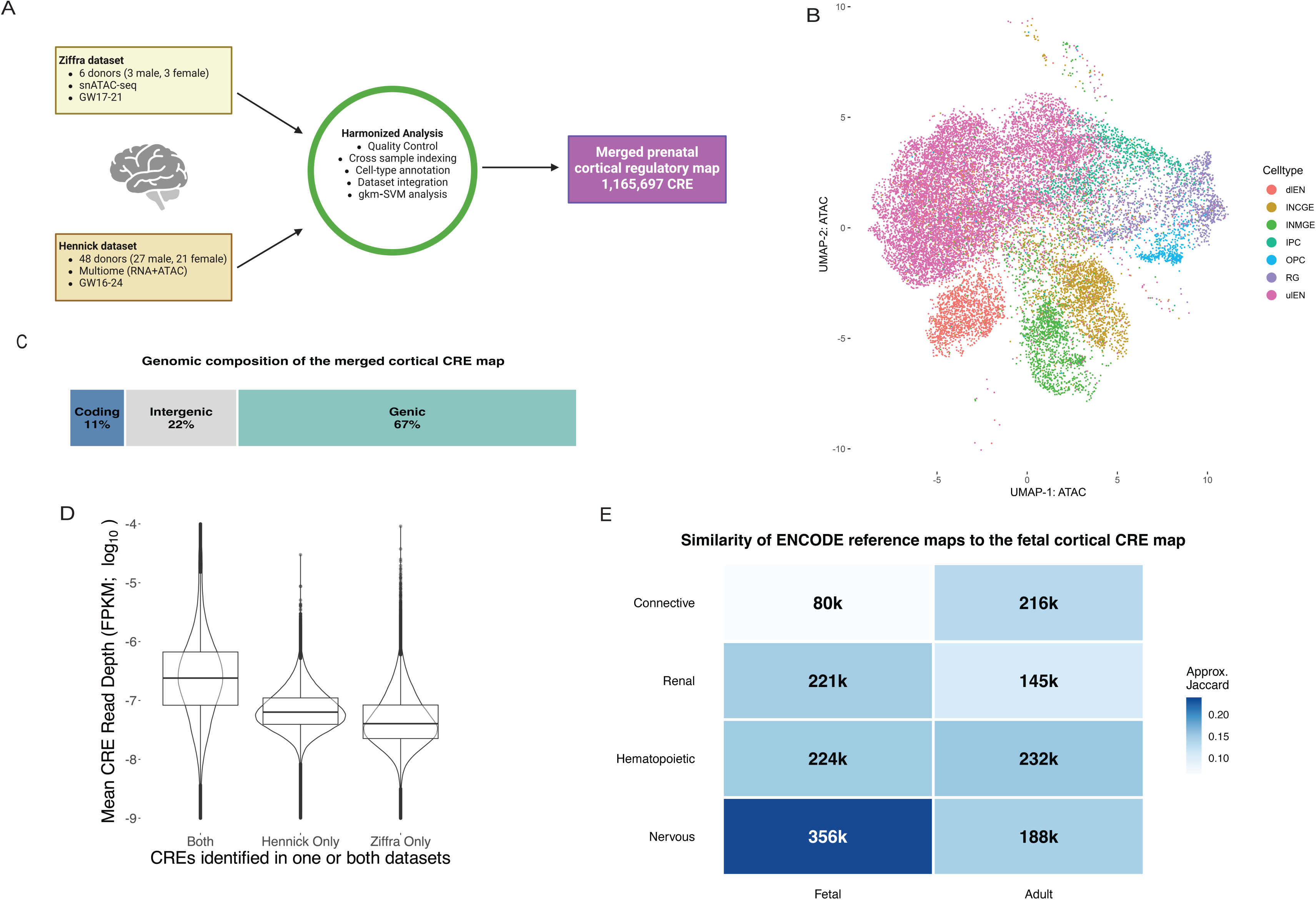
Construction and validation of a unified prenatal cortical CRE reference map. **(A)** Study design and harmonized analytical workflow. Single-nucleus chromatin accessibility (snATAC-seq) data from the Ziffra^37^ fetal cortical dataset (6 donors; 3 male, 3 female) and the Hennick^39^ fetal cortical Multiome dataset (48 donors; 27 male, 21 female) were processed using a shared analysis framework including quality control, cross-sample CRE indexing, cell-type annotation, and dataset integration. These analyses yielded a merged prenatal cortical regulatory map comprising 1,165,697 CREs. **(B)** Annotated UMAP of fetal cortical cell classes used to organize the merged regulatory map, including deep-layer excitatory neurons (dEN), upper-layer excitatory neurons (ulEN), MGE-derived interneurons (INMGE), CGE-derived interneurons (INCGE), intermediate progenitor cells (IPC), oligodendrocyte progenitor cells (OPC), and radial glia (RG). **(C)** Genomic composition of the merged cortical CRE map. Of the 1,165,697 CREs, 11% overlapped coding sequences, 67% were genic (within 50kb of a coding gene), and 22% intergenic. **(D)** Accessibility support for CREs identified in one or both datasets. CREs shared between the Hennick and Ziffra maps showed higher mean accessibility signal than CREs unique to either dataset, identifying a core set of robustly accessible cortical regulatory elements. **(E)** Validation of the reference cortical CRE map by comparison to ENCODE reference tissues.^15^ Tile color indicates approximate interval-level Jaccard similarity between the fetal cortical CRE map and merged ENCODE tissue reference maps, and tile labels indicate the number of overlapping CREs. Similarity was greatest for nervous-system reference maps, particularly fetal nervous tissue.

We identified 410,278 (35.1%) CREs shared between the two datasets, representing a putative core set of high-confidence regulatory elements. We identified 410,278 (35.1%) CREs shared between the two datasets, corresponding to 53.4% of the Ziffra map and 58.4% of the Hennick map. Although the datasets differed substantially in donor number (6 versus 48), the Ziffra dataset contained more nuclei (66,001 versus 38,265) and identified more CREs (768,785 versus 702,326), indicating that donor number alone does not explain the observed overlap. These shared CREs exhibited consistently higher read depth than dataset-specific CREs, supporting their robustness across studies (Figure 1D). Of course, their higher read-depth may also signify their greater activity. As external validation, we compared our cortical map with regulatory maps generated from ENCODE fetal and adult tissues using the same analytical framework. The cortical map showed greatest similarity to nervous-system reference maps, with fetal nervous tissue displaying the highest concordance at 30.6% similarity (356,490 CREs), while hematopoietic, renal and connective fetal and adult tissue were 6.9-19.9% similar (79,918-231,518 CREs) (Figure 1E).

### Annotation and classification of high-confidence regulatory elements

Because repetitive sequence can obfuscate genomic assignment and downstream sequence-based interpretation, we first quantified repeat content across all CREs using RepeatMasker.^48^ Repeat content varied widely across elements, with a median of 34%; 38% of CREs contained no annotated repetitive sequence, whereas 23% were composed of >90% repeats. Therefore, we classified CREs as low (≤25%; 540,194 CREs), moderate (>25% to ≤75%; 249,811 CREs), or high (>75%; 375,692 CREs) repeat content based on their quartiles. To determine whether repeat burden was associated with genomic annotation, we compared repeat-content classes across coding, genic, and intergenic CREs. CREs overlapping coding sequence were strongly enriched for low-repeat content (77% low, 14% moderate, 9% high). In contrast, non-coding compartments showed broadly similar distributions: genic CREs were 42% low, 23% moderate, and 35% high repeat and intergenic CREs were 44%, 20%, and 36%, respectively (Figure 2A). Thus, outside coding regions, repeat burden was not strongly structured by CRE location. We further found that ambiguous mapping increased substantially with repeat content: among 10,000 CREs randomly sampled from each repeat-content class, the fraction of CREs with multiple genomic alignments was 3.3% for low-repeat CREs, 3.4% for moderate-repeat CREs, and 8.2% for high-repeat CREs (Figure 2B). We retained high-repeat-content CREs because repetitive sequence is a *bona fide* component of the developing cortical regulatory landscape, and excluding these elements *a priori* would risk removing biologically relevant accessible sites; instead, we preserved them in the atlas while explicitly annotating repeat burden and mapping ambiguity to support cautious interpretation.

**Figure 2.**
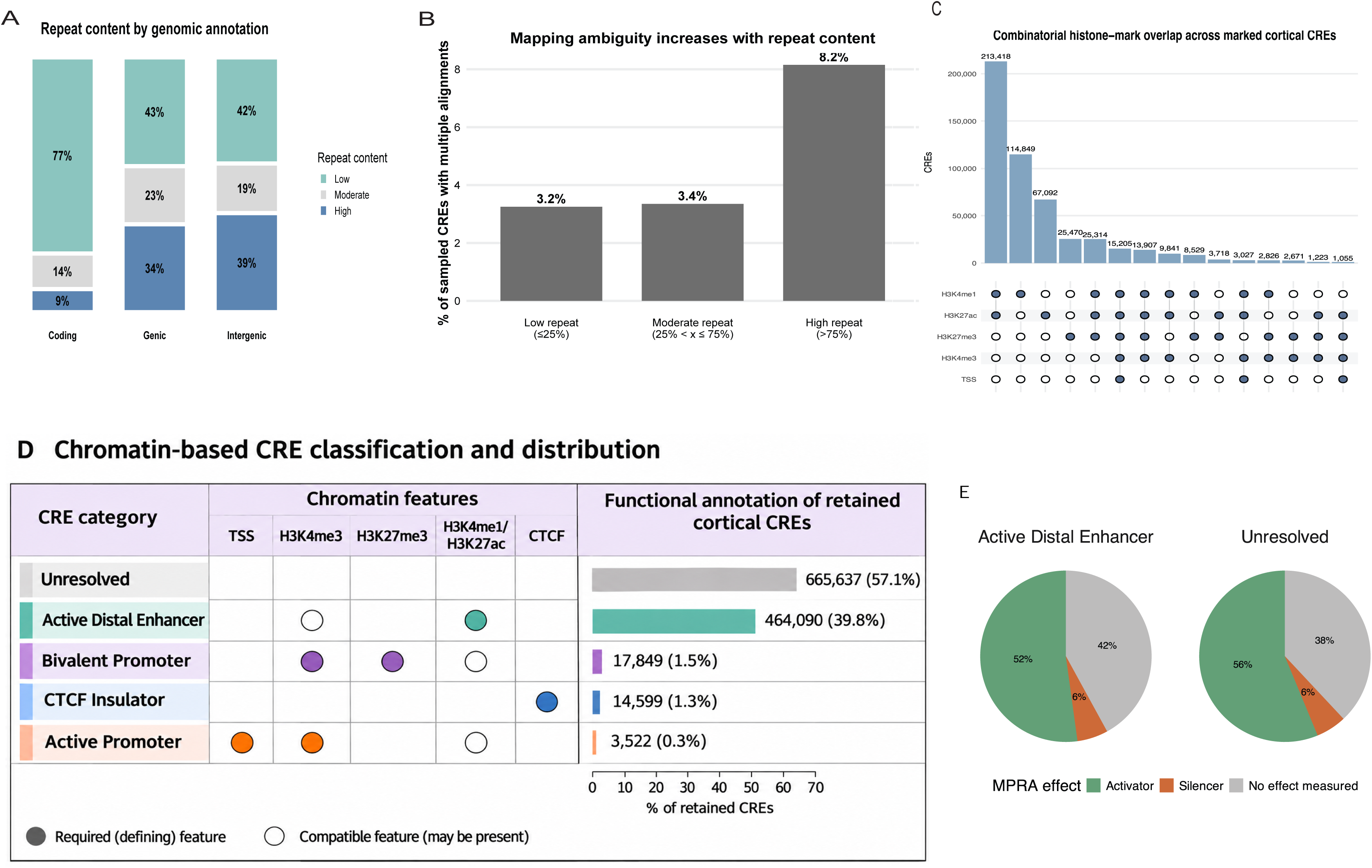
Repeat-aware thresholding and chromatin-based annotation of cortical CREs. **(A)** Repeat content by genomic annotation. CREs overlapping exons were strongly enriched for low-repeat sequence, whereas genic and intergenic non-coding CREs showed broadly similar repeat-class distributions. **(B)** Mapping ambiguity increases with repeat content. Among 10,000 CREs sampled from each repeat-content class, the fraction exhibiting multiple genomic alignments rose from 3.2% to 3.4% in low- and moderate-repeat CREs to 8.2% in high-repeat CREs. **(C)** Combinatorial overlap of chromatin features across marked cortical CREs. The most frequent combinations of TSS proximity (within 1 kb), H3K4me3, H3K4me1, H3K27ac, and H3K27me3 are shown; CTCF is used in the annotation schematic in panel D but is not displayed in this combination plot, illustrating layered and overlapping chromatin signatures among elements in the cortical map. **(D)**Chromatin-based classification and functional annotation of retained cortical CREs. Schematic showing the chromatin features used to classify CREs as active promoters, bivalent promoters, active distal enhancers, or CTCF-associated insulators. Filled circles indicate defining features, and open circles indicate compatible but non-defining features. CREs not assigned to these categories were classified as unresolved. Horizontal bars show the number and percentage of retained cortical CREs in each category, ordered from most to least abundant: unresolved (665,637; 57.1%), active distal enhancers (464,090; 39.8%), bivalent promoters (17,849; 1.5%), CTCF-associated insulators (14,599; 1.3%), and active promoters (3,522; 0.3%). **(E)** Functional outcomes among the subset of cortical CREs overlapping elements assayed in an independent published MPRA dataset. Pie plots show the proportions of tested histone-marked enhancers and unresolved CREs classified as activators, silencers, or having no detectable effect. The assayed subset comprised 36,037 active distal enhancers (7.8% of that class) and 9,737 unresolved CREs (1.5% of that class); these proportions should not be generalized to all cortical CREs.

Next, we functionally annotated the CREs using histone mark and architectural protein profiles by integrating publicly available ChIP-seq and CUT&Tag data for H3K27ac, H3K4me1, H3K4me3, H3K27me3, and CTCF across relevant tissues and cell types. In total, 45% of CREs (528,309 CRE) overlapped at least one chromatin mark and combinatorial signatures were common, consistent with layered regulation in the developing cortex (Figure 2E).^15,49,50^ We therefore used chromatin context to define operational and interpretable categories of CREs (Figure 2C): active promoters (H3K4me3 ± H3K27ac within ±1 kb of the transcriptional start site (TSS)), bivalent promoters (H3K4me3 + H3K27me3), active distal enhancers (either H3K4me1 or H3K27ac), and CTCF-associated insulators.^35,51–56^ Promoter-proximal CREs showed substantial bivalency: 84% carried both H3K4me3 and H3K27me3, a chromatin signature previously associated with transcriptional poising in neuronal progenitor states.^53,54^ Applying these criteria, we classified 3,522 active promoters (0.3%), 17,849 bivalent promoters (1.5%), 464,090 active distal enhancers (39.8%), and 14,599 CTCF-associated insulators (1.3%); 665,637 CREs (57.1%) remained unresolved under these chromatin annotations (Figure 2D). We anticipate that expanding cell type-matched chromatin annotations, for example through snCut&Tag,^57^ may resolve more of these sites in future studies.

To assess whether these annotation classes differed from expectations, we intersected these cortical CREs with results from an independent, previously published massively parallel reporter assay (MPRA) of 102,767 CREs on primary human mid-gestation cortical cells.^58^ Importantly, only 46,164 CREs in the cortical map overlapped elements assayed in that study, corresponding roughly to the 46,370 sequences in the MPRA determined to be accessible in primary cortical tissue and passing quality control. Among active distal enhancers, 36,037 CREs were tested (7.8% of active distal enhancers), of which 18,802 (52%) showed activator activity, 2,091 (6%) silencer activity, and 15,144 (42%) had no detectable effect; the remaining 428,053 enhancer CREs were not assayed. Surprisingly, among unresolved CREs, 9,737 overlapped tested MPRA elements (1.5% of unresolved CREs), of which 5,477 (56%) functioned as activators, 557 (6%) as silencers, and 3,703 (38%) showed no detectable effect (Fig. 2F); 655,900 unresolved CREs were not represented in the external MPRA dataset. Thus, aAlthough only a minority of cortical CREs could be evaluated through overlap with the cortical MPRA study,^58^ the assayed subset indicates that both histone-marked enhancers and unresolved CREs show regulatory activity. Notably, unresolved CREs were at least as likely as histone-marked enhancers to show activator activity in reporter assays, despite lacking canonical chromatin signatures. These results suggest that the unresolved category contains many*bona fide* regulatory elements that are not classified by the available histone-mark-based annotations. Perhaps sequence variation within these CREs and their contribution to human phenotypes can further clarify their genetic role.

### Genomic distribution of CREs

We next examined the genomic distribution of cortical CREs across chromosomes and functional classes. At the chromosome level, total CRE abundance was strongly associated with available non-repeat mapped sequences, i.e. CRE abundance was directly correlated with chromosome size (Supplemental Figure S1). Sex chromosomes, however, contained fewer CREs relative to their size. Functional classes differed, however, in their relationship to gene abundance. Promoter-proximal CRE counts tracked chromosome-level gene number very closely (Pearson r = 0.99; Spearman ρ = 0.98), whereas enhancer, insulator, and unresolved CRE counts showed substantial yet weaker associations (enhancer: Pearson r = 0.74, Spearman ρ = 0.65; insulator: Pearson r = 0.82, Spearman ρ = 0.77; unresolved: Pearson r = 0.69, Spearman ρ = 0.65) (Figure S2). These results indicate that broad chromosome-level CRE abundance is largely constrained by available genomic territory, while promoter-proximal regulatory elements are more tightly coupled to gene-rich genomic regions than other CRE classes.

### Distribution of CREs across cortical lineages reveals lineage-specific regulatory organization

We constructed cell-type-specific CRE maps within a common genomic indexing framework and added unified functional annotations and refined transcription factor (TF) motif analysis. To increase power and interpretability, we pseudo-bulked nuclei into major cortical lineages (“cell types”)—radial glia (RG), intermediate progenitor cells (IPCs), deep-layer excitatory neurons (dlEN), upper-layer excitatory neurons (ulEN), MGE-derived interneurons, CGE-derived interneurons, and oligodendrocyte precursor cells (OPCs)—based on linked gene expression or previously published annotations (Figure 1B). ^37,39^ Across lineages, we identified between 300,350 and 486,935 accessible CREs per lineage (Figure 3A). Upper-layer excitatory neurons harbored the largest CRE set (486,935 CREs from 55,753 nuclei), followed by deep-layer excitatory neurons (422,140 CREs from 11,410 nuclei) and radial glia (410,962 CREs from 7,675 nuclei), whereas interneuron lineages contained relatively fewer CREs relative to the number of nuclei collected (300,350 CREs from 10,715 CGE interneuron nuclei and 301,309 CREs from 9,858 MGE interneuron nuclei). Upper-layer excitatory neurons account for approximately half of all nuclei analyzed, suggesting that sampling depth was sufficient to approach saturation of CRE discovery in this cell population. Each lineage also contained a substantial set of lineage-specific CREs, ranging from 55,281 in CGE interneurons to 114,889 in OPCs. As a fraction of the lineage regulome, unique CREs comprised 18.4% of CGE interneuron CREs, 19.5% of MGE interneuron CREs, 19.6% of ulEN CREs, 23.2% of IPC CREs, 25.1% of RG CREs, 27.1% of dlEN CREs, and 34.5% of OPC CREs (Figure 3A). Thus, although all lineages retained a substantial shared regulatory component, lineage-restricted accessibility accounted for a substantial (∼18% - 35%) fraction of the cortical regulome, particularly in OPCs.

**Figure 3.**
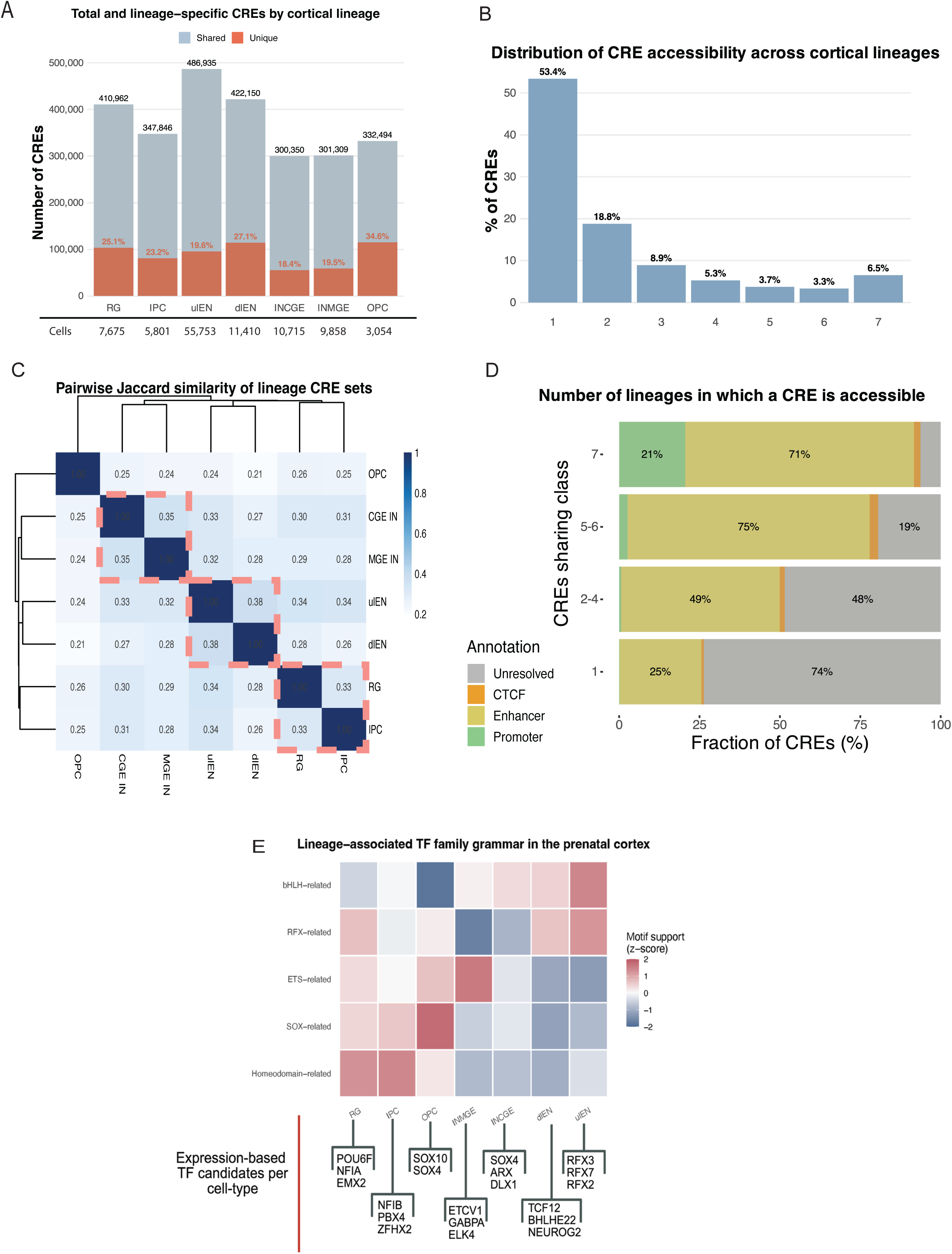
Lineage organization of chromatin accessibility in the developing human cortex. **(A)** Total and lineage-specific CREs identified in pseudo-bulked cortical lineages. Bars show total CREs for radial glia (RG), intermediate progenitor cells (IPCs), upper-layer excitatory neurons (ulEN), deep-layer excitatory neurons (dlEN), CGE-derived interneurons, MGE-derived interneurons, and oligodendrocyte precursor cells (OPCs). Orange segments indicate lineage-specific CREs, with percentages shown for each lineage. The number of cells harvested for each lineage is shown below each corresponding bar. **(B)** Distribution of CRE accessibility across cortical lineages. Bars show the percentage of CREs in the merged cortical reference map detected in 1 to 7 lineage pseudo-bulks. Most CREs are restricted to a single lineage, whereas only a small fraction is shared across all seven lineages. **(C)** Pairwise Jaccard similarity of lineage CRE sets. Similarity is highest between upper- and deep-layer excitatory neurons, between CGE- and MGE-derived interneurons, and between RG and IPCs, these three pairs being equally similar to one another, consistent with developmental relatedness among these populations (each pair is highlighted). OPCs are the most distinct lineage. **(D)** Functional composition of CREs stratified by lineage-sharing breadth. Horizontal stacked bars show the fraction of CREs in each sharing class (1, 2–4, 5–6, or 7 lineages) annotated as promoter, enhancer, CTCF-associated, or unresolved. Lineage-specific CREs are dominated by unresolved and enhancer annotations, whereas CREs shared across all seven lineages show the highest promoter fraction. **(E)** Family-level motif support across cortical lineages. deltaSVM-derived motif support scores were grouped into broad TF families and summarized across major cortical lineages. Values were z-score normalized within family across lineages to highlight relative lineage-associated motif support. Homeodomain-related motifs were strongest in RGs and IPCs, SOX-related motifs in OPCs, ETS-related motifs in MGE-derived interneurons, bHLH-related motifs in deep-layer excitatory neurons, and RFX-related motifs in upper-layer excitatory neurons. Representative family members were prioritized using lineage-matched TF gene expression.

We next asked how broadly individual CREs were shared across cortical lineages. Across the merged cortical CRE map, accessibility was most often restricted to a single lineage, with 53.4% of CREs detected in only one pseudo-bulked lineage (Figure 3B). In contrast, only 6.5% of CREs were accessible across all seven lineages. Cell-type overlap significantly deviated from an assumed Poisson distribution (p < 0.0001), with an enrichment of CREs specific to a single cell-type as well as an enrichment of CREs found in all cell-types. Intermediate sharing states were less common, comprising 18.8% of CREs in two lineages, 8.9% in three, 5.3% in four, 3.7% in five, and 3.3% in six lineages. These results indicate that the fetal cortical regulome is dominated (∼80%) by lineage-restricted (3 or fewer related lineages) elements with a much smaller core of broadly shared accessible sites.

Pairwise comparison of lineage CRE sets further showed that chromatin accessibility recapitulates developmental relatedness (Figure 3C). The greatest similarity was observed between deep- and upper-layer excitatory neurons (Jaccard index = 0.38), followed by CGE- and MGE-derived interneurons (0.35), with radial glia and IPCs also showing substantial similarity (0.33). More moderate similarities were observed among progenitor and excitatory neuronal lineages and between interneuron lineages and other cell classes. In contrast, OPCs were the most distinct lineage overall, with the pairwise Jaccard index ranging from 0.21 to 0.26, relative to the other groups. Together, these findings indicate that developmentally related cortical lineages share a larger fraction of their accessible regulome than more distant populations.

To determine whether lineage-sharing breadth corresponded to distinct functional classes of CREs, we stratified elements by the number of lineages in which they were accessible and re-examined their chromatin-state annotations (Figure 3D). CREs accessible in only a single lineage were composed predominantly of unresolved elements (74%), with active distal enhancers comprising 25% and promoters or CTCF-associated sites contributing even smaller fractions. By contrast, enhancers dominated CREs shared across 2–4 lineages (49%) and became even more prominent among CREs shared across 5–6 lineages (75%). CREs accessible in all seven lineages remained primarily enhancer annotated (71%) but showed the strongest promoter contribution (21%) of any sharing class. Together, these results indicate that lineage-specific and moderately shared cortical CREs are largely enhancer-rich, whereas the small set of pan-lineage CREs contains a comparatively greater promoter-proximal component. This organization supports a model in which cortical lineages share a common accessible regulatory backbone while lineage identity is implemented primarily through selective deployment of distal regulatory elements.

### Transcription factor sequence-grammars underly cortical lineage specification

Building on our cell type–resolved CRE maps, we next asked which transcription factor (TF) sequence features distinguished lineage-specific regulatory elements. We applied our prior machine learning algorithms^44–47^, using discriminative short-sequence feature (11-mer sequence grammars) distributions and mapping enriched sequence featuresed to known TF motifs and motif families,^43,45^ to the lineage-resolved CRE sets. Across the lineage-resolved maps, 149 motif-associated TF features were both enriched at least 4-fold for the top 5% of 11-mers and expressed in the top 50% of genes within a celltype in at least one cell type, spanning diverse families including homeodomain, bHLH, SOX-related, RFX-related, POU-domain, ETS-related, and nuclear factor families (Figure 3E; Supplementary Table 1). At the broadest level, progenitor-associated lineages were enriched for homeodomain-related sequence features, whereas neuronal lineages showed stronger neurogenic bHLH-associated signatures. More specifically, RG and IPC CREs were enriched for homeodomain-, paired-, and HD-LIM-related sequence features, consistent with regulatory programs linked to positional patterning, cortical arealization, and maintenance of early developmental competence.^59,60^ By contrast, neuronal lineages showed stronger bHLH-associated sequence features, likely reflecting neurogenic and differentiation-associated programs linked to canonical regulators such as NEUROG- and NEUROD-related factors.^61,62^ This broad homeodomain-to-bHLH transition matches the expected arc of corticogenesis, in which spatially patterned progenitor states give way to transcriptional programs that drive neuronal commitment, subtype specification, and maturation. Family-level enrichment among TFs with strongest lineage-associated support further reinforced this developmental logic, including significant enrichment of homeodomain-related factors in IPCs (odds ratio, OR = 3.21, false discovery rate, FDR = 0.013) and radial glia (OR = 3.68, FDR = 0.024), ETS-related factors in MGE-derived interneurons (OR = 154.12, FDR = 1.8 × 10^-8^), bHLH-related factors in deep-layer excitatory neurons (OR = 19.62, FDR = 3.5 × 10^-5^), and RFX-related factors in upper-layer excitatory neurons (OR = 19.05, FDR = 0.035).

Because motifs often reflect family-level binding preferences rather than activity of a single TF, we next used lineage-matched expression to prioritize representative family members within these enriched programs.^39^ This approach prioritized distinct candidate regulators across the major cortical lineages (Figure 3E, Supplemental Table 2). CGE-derived interneurons were associated with SOX4-, ARX-, and DLX1-related signatures, whereas MGE-derived interneurons highlighted TCF12- and POU2F1/2-associated programs. Progenitor-associated populations recovered NFIA, POU6F1, and EMX2 in radial glia and NFIB, ZFHX2, and PBX4 in IPCs, consistent with related but distinguishable progenitor-state programs involved in cortical patterning and progenitor differentiation.^59,63,64^ OPCs showed the expected enrichment of SOX10, together with SOX4, supporting a glial-lineage-associated regulatory grammar distinct from neuronal programs.^65,66^ Excitatory neuron lineages were likewise associated with discrete candidate modules, including TCF12, BHLHE22 and NEUROG2 in deep-layer excitatory neurons and RFX family members (RFX3, RFX7, RFX2) in upper-layer excitatory neurons.^67,68^ Notably, the prominence of RFX3 is consistent with a recent study implicating altered RFX3-dependent regulatory networks in language-associated cortex in ASD, particularly in glutamatergic neuronal populations, providing an external point of convergence for our findings.^69^ Together, these results indicate that lineage-specific cortical CREs encode distinct and biologically interpretable TF sequence grammars that can be used to prioritize representative candidate regulators when integrated with lineage-matched TF expression.

### CRE–gene linkage reveals predominantly local but multi-gene regulatory architecture

We next asked which genes might be associated with prenatal cortical CREs through predicted chromatin contacts. Using a maximum distance of 500 kilobase, we identified 5.44 million CRE–gene links involving 983,768 CREs and 11,800 gene transcription start sites across the merged prenatal cortical atlas. We then used C.Origami,^70^ a multimodal deep neural network algorithm that performs *de novo* prediction of cell-type-specific chromatin organization using DNA sequence and cell-type-specific genomic features to predict CRE-gene contact frequencies. In most cases, the strongest linked gene corresponded to the nearest gene’s TSS (87.6%), whereas the remaining (12.4%) CREs showed stronger linkage to a more distal target. Overall, 175,115 CREs (17.8%) linked to a single gene, while 379,363 CREs (38.6%) linked to between 2–4 genes (89.6% of these being most strongly linked to the nearest gene), 283,219 CREs (28.8%) to 5–9 genes (nearest gene strongest for 84.7%), 119,117 CREs (12.1%) to 10–19 genes (nearest gene strongest for 76.5%), and 26,954 (2.7%) to 20 or more genes (nearest gene strongest for 60% of CREs). As the number of genes near a given CRE increased, so did the apparent likelihood of it having a stronger predicted contact with a distal gene, however nearly all increase in interaction frequency lay withinone standard deviation (chromatin contact scores were grouped by distance into 8,192 base-pair buckets, corresponding to cOrigami units; mean and standard deviation were each bucket; 88% of measured CRE-TSS contact scores were within 1 standard deviation of the distance bucket), indicating that prenatal cortical regulatory architecture is predominantly local but not reducible to simple nearest-gene assignment based on available data.

Linkage structure also differed across annotation classes. Active distal enhancers and unresolved CREs each linked to a median of 4 genes per element, whereas active and bivalent promoters linked to a median of 6 and 7 genes, respectively. Promoter-associated CREs were also less likely than enhancers or unresolved elements to have the nearest gene as their strongest linked target, suggesting that promoter-rich elements may participate in more complex local regulatory neighborhoods. Linkage coverage was likewise not uniform across CRE classes, with promoter-associated CREs most readily resolved by the model and unresolved CREs least readily resolved. Together, these results indicate that candidate target-gene architecture differs systematically across functional classes of prenatal cortical CREs although elucidating their specific genomic properties will require more work.

### Inter-donor recurrence, developmental persistence, and sex bias identify the stable and variable compartments of the cortical regulome

We next asked whether additional axes of biological variation—across individuals, developmental time, and sex—follow the same core organizational logic observed across all cortical lineages. Donor-level recurrence was heavily skewed toward low-frequency elements, indicating that the majority of cortical CREs are detected in only a subset of individuals, all of whom differ in genomic sequence. Across the 54-donor atlas, 383,370 CREs (32.9%) were not recovered in any donor-specific map under current thresholds (and, thus, detectable only in the aggregate), 250,022 (21.4%) were detected in only a single donor, and 262,491 (22.5%) were detected in 2–4 donors. By contrast, only 42 CREs (0.0036%) were detected across all 54 donors. Among CREs detected in at least one donor, recurrence was strongly associated with normalized accessibility strength, with median baseMean increasing progressively from 0.35 in single-donor CREs to 0.49, 0.85, 1.48, 3.54, 22.6 and 99.4 in CREs detected in 2–4, 5–9, 10–19, 20–39, 40–53 and all donors, respectively (Figure S2). Recurrence classes also differed markedly in annotation: low-recurrence CREs were predominantly unresolved, intermediate-recurrence CREs became increasingly enhancer-dominant, and the most recurrent CREs showed the strongest promoter-proximal and bivalent contribution. Thus, inter-individual recurrence reveals a large weak and variable compartment together with a very small, strong, promoter-rich core.

A similar pattern was evident across developmental time. CREs detected across different numbers of gestational weeks showed distinct annotation compositions, with single-week CREs enriched for unresolved elements, intermediate persistence classes becoming progressively enhancer-dominant, and CREs detected across all nine gestational weeks showing the strongest promoter-proximal and bivalent contribution. Normalized accessibility strength also increased steadily with developmental persistence, with median baseMean rising from 0.51 in CREs detected in only one gestational week to 0.80, 2.19 and 15.80 across 2–4, 5–8 weeks and all nine gestational weeks (Figure S3). Thus, developmental persistence recapitulates the same regulatory architecture observed across lineages and donors: restricted CREs are weak and enriched for unresolved annotations, whereas broadly persistent CREs are stronger and enriched for promoter-associated states.

Importantly, a limitation of this test is that a significant portion of CREs were present in the merged consensus atlas but did not meet peak-calling thresholds in donor- or age-stratified maps. This could reflect the underlying biology but surely is affected by the lower statistical power of individual versus global maps, as well as the uneven distribution of celltypes that contribute to donor- and age-stratified maps. Additionally, the stable and broadly shared CRE compartments identified across donors, gestational persistence, and lineages were also the classes most readily resolved by the CRE-linkage model, whereas the variable unresolved compartment remained less well resolved at the level of target-gene assignment.

### Sex-biased CREs comprise a small but distinct component of the cortical regulome

We finally tested whether chromatin accessibility differed systematically between the male and female cortex, given the pervasive sex-differences in neurodevelopmental disorders. Threshold-based overlap of sex-stratified CRE maps indicated that 523,396 CREs were detected in both sexes, whereas 382,683 and 259,618 were detected only in female- (24 donors) and male- (30 donors) aggregated maps, respectively. However, statistical analysis identified relatively few that were significantly differentially accessible: 1,048 were male-biased and 98 were female-biased, compared with 1,164,551 unbiased CREs (Figure 4A). Thus, although threshold-based CRE recovery differed substantially between sex-aggregated maps, quantitative sex-associated differences in accessibility were limited to a very small subset of the cortical regulome.

**Figure 4.**
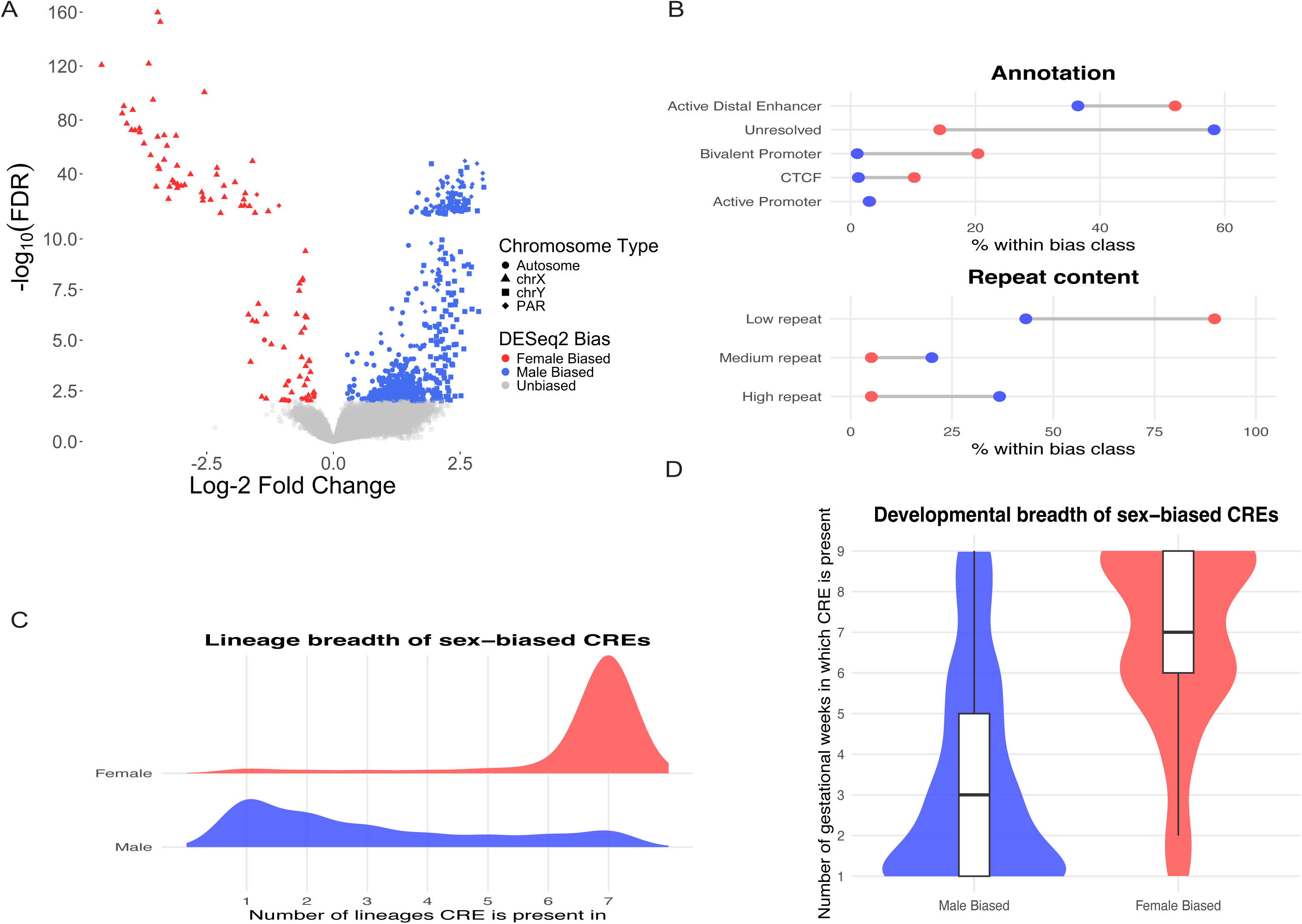
Sex-biased CREs are rare overall but differ in annotation, repeat content, and lineage breadth. **(A)** Volcano plot of differential chromatin accessibility between male and female fetal cortex. Most cortical CREs were not significantly sex-biased: only 1,048 male-biased and 98 female-biased CREs were identified. Points are colored by bias direction and chromosome class, distinguishing autosomal from sex chromosome-linked CREs. **(B)** Functional annotation and repeat-content composition of sex-biased CREs. Female-biased CREs were enriched for active distal enhancers and bivalent promoters and had predominantly low repeat content, whereas male-biased CREs were dominated by unresolved elements and showed substantially greater repeat content. Values indicate the percentage of CREs within each bias class. **(C)** Lineage breadth of sex-biased CREs across seven major cortical lineages. Female-biased CREs were concentrated among broadly shared elements present across all seven lineages, whereas male-biased CREs were more often restricted to a smaller number of lineages. **(D)** Developmental breadth of sex-biased CREs across gestational weeks. Female-biased CREs were detected across more developmental time points overall, whereas male-biased CREs were typically restricted to fewer gestational weeks, indicating greater temporal persistence of the female-biased CRE subset.

Sex-biased CREs were strongly partitioned by chromosome class (Figure 4A). Female-biased CREs were overwhelmingly X-linked, with 90 of 98 (91.8%) mapping to the sex-linked portion of the X chromosome (chrX), 2 (2.0%) to the pseudo-autosomal portion of chrX and only 6 (6.1%) on autosomes. In contrast, male-biased CREs were distributed primarily across autosomes (789 CREs, 75.3%) and the Y chromosome (chrY) (181 CREs, 17.3%), with only 78 CREs (7.4%) mapping to pseudo-autosomal regions; only two elements mapped to chrX. Thus, female-biased accessibility was largely driven by X-linked elements, whereas male-biased accessibility was dominated by autosomal and Y-linked CREs.

Sex-biased CREs differ in their functional annotation profiles as well (Figure 4B). Female-biased CREs were composed primarily of active distal enhancers (51 of 98; 52.0%), with additional contributions from bivalent promoters (20 of 98; 20.4%), unresolved CREs (14 of 98; 14.3%), and CTCF-associated elements (10 of 98; 10.2%), whereas active promoters were rare (3 of 98; 3.1%). In contrast, male-biased CREs were dominated by unresolved elements (611 of 1,048; 58.3%), followed by active distal enhancers (382 of 1,048; 36.5%), while promoter- and CTCF-associated classes each contributed only small fractions. Female biased promoters (on chrX) tended to be near genes that escape X-inactivation (*XIST, SYAP1, EIF2S3, TXLNG, ZFX, DDX3X, AP1S2, STS, ZRSR2, CDK16, USP9X, KDM6A, TRAPPC2, RPS4X, KDM5C*), and were generally correlated with female biased expression, whereas genes near male biased promoters were not correlated with biased expression (excluding chrY genes). Repeat content also differed markedly between the two classes (Figure 4B). Female-biased CREs were overwhelmingly low repeat (90.8%), with only 4% and 5.1% in medium- and high-repeat sequence respectively. Male-biased CREs were substantially more repeat-rich, with 43.2% low-, 20.1% medium-, and 36.6% high-repeat sequence (Figure 4B). Together, these results indicate that male- and female-biased accessibility differences arise in distinct chromatin and sequence compartments.

These differences were mirrored in developmental persistence and lineage sharing (Figures 4C, 4D). Female-biased CREs were broadly persistent across gestational weeks, with a mean of 6.55 gestational weeks per CRE, and were also broadly distributed across cortical lineages, with a mean of 6.46 lineages and a median of 7. In contrast, male-biased CREs were substantially narrower in both dimensions, with a mean of 2.84 gestational weeks and 3.06 lineages per CRE, and a median lineage distribution of only 2. Consistent with this broader persistence, female-biased CREs also showed substantially higher normalized accessibility than male-biased or unbiased CREs (median baseMean 7.91, 1.04, and 0.398, respectively; Figure S3). Donor recurrence further distinguished the two classes: female-biased CREs were concentrated in highly recurrent donor classes, whereas male-biased CREs were most often detected in low- to intermediate-recurrence groups. Thus, female-biased CREs define a small, strong, and broadly shared regulatory compartment, whereas male-biased CREs occupy a larger but more heterogeneous and restricted component of the cortical regulome.

Finally, we asked whether male-biased CREs were preferentially associated with specific cortical lineages. Because individual CREs can be accessible in multiple lineages, we first assessed overlap enrichment relative to unbiased CREs. Male-biased CREs were most strongly enriched among CREs accessible in upper-layer excitatory neurons (OR = 2.94, FDR = 4.6 × 10^-64^), followed by deep-layer excitatory neurons (OR = 2.00, FDR = 2.7 × 10^-68^), radial glia (OR = 1.74, FDR = 1.4 × 10^-18^), and MGE-derived interneurons (OR = 1.70, FDR = 9.9 × 10^-16^), with weaker but still significant enrichment in CGE-derived interneurons and IPCs; OPC overlap was not enriched (OR = 0.96, FDR = 0.61). To simplify interpretation, we next restricted analysis to the subset of male-biased CREs present in only a single lineage. Among these 332 lineage-specific male-biased CREs, the largest fraction was observed in upper-layer excitatory neurons (32.2%), followed by radial glia (22.0%) and deep-layer excitatory neurons (17.8%), whereas smaller fractions were assigned to CGE-derived interneurons (7.8%), MGE-derived interneurons (7.5%), OPCs (7.2%), and IPCs (5.4%). Together, these results indicate that sex-associated chromatin differences are rare overall, but male- and female-biased CREs occupy sharply distinct chromosomal, regulatory, and developmental regimes, with the lineage-restricted component of male-biased accessibility concentrated primarily in excitatory neuronal and radial glial regulatory programs.

## Discussion

A central finding of this study is the reframing of the prenatal cortical regulome not as a static catalog of functional elements but as a hierarchically organized system with stable and variable regulatory compartments. Most chromatin accessibility studies largely focus on identifying cell-type-specific CREs, assigning them to known chromatin states, and connecting them to candidate disease loci.^28,37,71^ These goals remain essential, but they do not by themselves explain how regulatory variation is distributed or how it can be understood across developmental cell types or individuals. By integrating annotation, lineage sharing, developmental persistence, donor frequency, sex bias, TF sequence grammar, and CRE-gene linkage within a single regulatory framework, we identify a consistent organizational principle: a small strong, recurrent, promoter-rich core coexists with a much larger weak and variable compartment enriched for enhancer-like and currently unresolved elements. This significant pattern, rather than any single atlas component alone, is the main conceptual contribution of the study.

One implication of this framework is that the most dynamic part of the prenatal cortical regulome is also the part least well captured by canonical chromatin-state annotation.^72,73^ It is also the part most likely to explain human phenotypic variation. Across lineages, gestational persistence, donor recurrence, and sex bias, the variable compartment was repeatedly enriched for unresolved CREs. This observation would be less meaningful if unresolved elements were simply low-confidence leftovers, but our overlap with an external dataset argues against that interpretation. Many unresolved CREs were functionally active in reporter assays, and in some comparisons, they were at least as likely as histone-marked enhancers to show activator activity. Taken together, these findings suggest that unresolved CREs are not merely experimental noise but are instead a large and biologically important fraction of the prenatal cortical regulatory landscape. Indeed, many of them may be facilitators as described by Kassouf and colleagues.^36^ More broadly, our analyses suggests that existing histone-mark frameworks capture the stable backbone of prenatal cortical regulation more effectively than its variable component.^72,73^

What remains unclear is what biological classes are contained within the unresolved CRE compartment. Our data suggest that it is unlikely to represent a single category. Instead, unresolved CREs probably comprise a heterogeneous mixture that includes bona fide distal regulatory elements that were not captured in histone occupancy assays or were lacking matched chromatin-state information in the relevant prenatal cortical cell types, weak or transient elements that fall below the detection thresholds of current histone-mark frameworks, repeat-rich or otherwise technically difficult loci, and potentially non-canonical regulatory sequences whose activity is not well described by standard promoter/enhancer categories.^36^ Several observations argue that this class is not simply an aggregate of low-confidence sites. Unresolved CREs were consistently enriched among lineage-restricted, donor-restricted, and developmentally restricted accessibility classes, and many of the assayed subset showed reporter activity in an independent study despite lacking the chromatin signatures used for classification. Biologically, these features suggest that the unresolved compartment may be enriched for highly context-dependent developmental regulatory elements, including transient enhancers, lineage-selective distal elements, and other functional sites whose activity is not accompanied by canonical active histone marks, as has been observed for developmentally regulated loci more broadly and for some developmental enhancers *in vivo*.^74,75^ In that sense, unresolved CREs may represent not merely an annotation gap, but a dynamic component of the prenatal cortical regulome that is especially sensitive to cell state, developmental timing, and inter-individual variation. Indeed, genetic variation studies of these sequences in neurodevelopmental disease versus controls should shed light on their importance.

This view helps explain why lineage specificity in the prenatal cortex is concentrated in enhancer-rich and unresolved compartments rather than in promoter-associated elements. Promoter-rich CREs are disproportionately associated with strong signal, and recurrence across lineages, developmental windows, and donors. By contrast, lineage-restricted and developmentally restricted CREs show lower activity, on average, and are enriched for unresolved or distal regulatory annotations. This organization is biologically plausible. Promoter-associated elements are expected to anchor broadly required transcriptional programs, whereas distal elements are more likely to encode conditional regulatory logic that differs across cell states, developmental transitions, and individuals based on their sequence variation.^35,75,76^ In that sense, the prenatal cortex is regulated by a relatively small stable core layered beneath a much larger context-sensitive regulatory compartment.

Our TF sequence analysis reinforces this distinction. At the TF family level, progenitor-associated CREs showed a stronger homeodomain-related sequence grammar, whereas neuronal lineages are enriched for bHLH-associated signatures, recapitulating the expected shift from positional specification and developmental competence to neurogenic commitment and neuronal differentiation.^59,61–63^ More importantly, integrating sequence features with lineage-matched TF expression allowed us to move beyond broad family labels toward representative candidate regulators for specific cortical lineages. These candidates are unlikely to represent complete or exclusive regulatory programs, but they provide a set of tractable hypotheses for future perturbation studies. The recurrence of factors such as SOX10, ARX, DLX1, NFIA, EMX2, and RFX3 across distinct lineage contexts suggests that the sequence grammar of prenatal cortical CREs is sufficiently structured to recover known biology while also nominating candidate regulatory modules not obvious from accessibility alone.^63–65,77^ The prominence of RFX3 is particularly interesting in this regard. Our data identify RFX3 as a sequence- and expression-supported candidate in upper-layer excitatory neurons, pointing to a possible role in excitatory lineage regulatory architecture during prenatal development. The recent implication of altered RFX3-dependent networks in the language-associated cortex in ASD suggests that TF-centered regulatory modules identified in prenatal chromatin maps may remain relevant to later disease-associated molecular phenotypes.^69^ We do not interpret this as a direct replication across stages or tissues, but rather as evidence that prenatal regulatory grammars may help define the developmental substrates through which later disease-relevant dysregulation emerges.

The donor recurrence analysis gives a particularly important perspective on what an atlas can and cannot tell us. Most CREs were detected in only a subset of individuals, whereas only a tiny fraction was present across all donors. Some of this long low-recurrence tail is undoubtedly influenced by signal strength and detection threshold sensitivity; weak elements are harder to recover reproducibly across multiple experiments, such as donor-stratified maps. But the recurrence analysis was informative precisely because annotation and normalized accessibility changed systematically with recurrence class. Low-recurrence CREs were unresolved-dominant, intermediate-recurrence CREs were enhancer-rich, and the most recurrent compartment was promoter- and bivalent-enriched. This suggests that the donor-restricted component is not simply technical debris, but a plausible substrate for structured inter-individual regulatory variation.^17^ That interpretation is particularly relevant for complex trait genetics: the regulatory variation most likely to differ across individuals may reside in the least recurrent, least canonically annotated portion of the prenatal cortical CRE map.^1,4^

The developmental persistence and sex-bias analyses fit naturally into this same framework. CREs detected across many gestational weeks were stronger and increasingly promoter-associated, whereas temporally restricted CREs were weaker and more unresolved. Likewise, sex-biased CREs did not represent a broad reorganization of the prenatal cortical regulome but instead occupied specific and highly structured compartments. Female-biased CREs defined a small, strong, X-linked, recurrent, and developmentally persistent class enriched for annotated enhancer and bivalent promoter states. Male-biased CREs were more numerous but more heterogeneous, more unresolved, and more restricted across lineages and developmental time. The important point is not simply that sex differences exist, but that sex, like donor recurrence and developmental persistence, exposes the same underlying architecture of stable versus variable regulatory compartments.

The CRE-gene linkage analysis adds an important but still preliminary mechanistic layer. Most CREs were most strongly linked to the nearest gene, but not all, and most were linked to more than one candidate target. This is consistent with a regulatory architecture that is predominantly local but not reducible to nearest-gene assignment alone.^73,78,79^ What remains unresolved is whether the stable and variable CRE compartments differ systematically in target-gene architecture, pathway convergence, or network topology. That is likely to be an important direction for future work. If donor-restricted, developmentally restricted, or unresolved CREs preferentially connect to classes of genes or regulatory modules, that would provide a much stronger bridge between chromatin architecture and disease-relevant GRNs.

Several limitations of the present study follow directly from these conclusions. First, the large unresolved compartment almost certainly reflects, in part, incomplete cell type- and stage-matched chromatin-state annotation in the prenatal cortex. Second, our donor and gestational persistence analyses remain threshold-dependent and should not be interpreted as direct measures of absolute biological absence. Third, although the donor-restricted compartment is a plausible substrate for rare genotype-associated regulatory variation, systematic dissection of genetic background was beyond the scope of this study. Finally, CRE-gene linkage and TF sequence grammar identify candidate regulatory relationships rather than direct causal interactions and will require functional perturbation for validation.

Despite these limitations, the present work provides a useful framework for interpreting non-coding variation in the developing human cortex or in other tissues. Rather than asking only where accessible elements exist, this study asks how they are organized, shared, and vary across developmental and individual contexts. The answer is a prenatal cortical regulome built from a small stable core and a much larger variable component. For neurodevelopmental disorders such as ASD, this distinction may be especially important. It suggests that disease-associated non-coding variation may often reside not only in well-annotated promoters and enhancers, but also in weak, context-dependent, and incompletely annotated CREs that vary across lineages, developmental windows, and individuals.^18,19,22,26^ In that sense, the major contribution of this study is not simply a larger prenatal cortical reference maps and atlas, but a regulatory architecture for thinking about how non-coding variation acts in development.

## MATERIALS AND METHODS

### Data sources and preprocessing

Single-nucleus paired ATAC-seq and RNA-seq data were obtained from two prenatal cortical datasets: newly generated Multiome data on 48 mid-gestation prenatal cortices,^39^ and previously published single nucleus (sn)ATAC-seq data from 6 prenatal cortices,^37^ for a total of 54 donors and 104,266 nuclei. Reads were aligned to the Cell Ranger ARC 2.0.2 reference genome (GRCh38, GENCODE v32/Ensembl98) using cellranger-arc count. FASTQ files were additionally aligned to the CHM13v2.0 genome^80^ for sex-chromosome re-mapping and pseudo-autosomal region assignment. Mapped single-nucleus ATAC-seq reads were split by cellular annotation using Sinto (https://github.com/timoast/sinto) and pooled into pseudo-bulk datasets using samtools.^81^ Pseudo-bulked data were generated separately within each study using available cell-type labels and donor sex: 36 sex & cell-type pseudobulks were generated from the two datasets (male and female pseudobulks for each of 8 cell-types from the Ziffra dataset^37^ and 10 cell-types from the Hennick dataset^39^). For the Ziffra dataset,^37^ pseudo-bulks were generated for radial glia, intermediate progenitor cells, astrocyte/oligodendrocyte precursor cells, CGE-derived interneurons, MGE-derived interneurons, deep-layer excitatory neurons, upper-layer excitatory neurons, and early excitatory neurons. For the Hennick dataset,^39^ pseudo-bulks were generated for radial glia, dividing radial glia, intermediate progenitor cells, oligodendrocyte precursor cells, CGE-derived interneurons, MGE-derived interneurons, layer 2/3 excitatory neurons, layer 4 excitatory neurons, layer 5 excitatory neurons, and layer 6/subplate excitatory neurons. These cell type annotations were obtained from published data.^37,39^ Microglial and endothelial cells were excluded from downstream pseudo-bulking and analysis. Additional pseudo-bulked datasets were generated across all 54 donors and across gestational age groups (GW16– GW24). Donor-by-cell-type pseudo-bulks were also generated for read-counting and differential analyses.

### Peak-calling and reference CRE map generation

Peak calling was performed on each pseudo-bulk using MACS2^82^ with the parameters -- SPMR -q 0.01 --nomodel --ext-size 50 --shift 100. Peaks on scaffolds were excluded, retaining only chromosomes 1–22, X, and Y. Consensus CRE maps were then generated using the index method of Meuleman et al. (2020).^40^ MACS2 narrowPeak and summit outputs were converted into the required BED-like input format and compressed using BEDOPS starch. These starch files were used as input to the Index script to construct non-redundant consensus CRE maps. Separate reference maps were generated for each source dataset, and a merged prenatal cortical map was generated by integrating the two studies. Cell-type specific reference maps were generated by combining pseudo-bulks from matched or related annotations across studies as follows: (1) radial glia (RG), (2) intermediate progenitor cells (IPC), (3) oligodendrocyte precursor cells (OPC), (4) CGE-derived interneurons (IN-CGE), (5) MGE-derived interneurons (IN-MGE), (6) upper-layer excitatory neurons (ulEN), and (7) deep-layer excitatory neurons (dlEN). Cell-type contributions to the merged cortical map were assigned by overlap of pseudo-bulk peak calls with consensus CREs. Sex-chromosome and pseudo-autosomal assignments were refined using CHM13v2.0 re-mapping. CREs whose constituent reads re-mapped to the alternate sex chromosome with high concordance, or which lay within pseudo-autosomal intervals, were reclassified as pseudoautosomal (PAR) where appropriate.

### Functional annotation of CREs

Functional annotation of CREs was performed by intersecting the merged cortical CRE map with ENCODE ChIP-seq datasets and prenatal cortical CUT&Tag datasets from Hennick et al. (2025),^39^ including H3K27ac, H3K4me1, H3K4me3, H3K27me3, and CTCF. Overlap with transcription start sites (TSSs) was determined using UCSC knownGene annotations for coding genes; when gene annotations had multiple transcripts and in turn multiple TSSs, the most upstream TSS was used. CREs were classified using an operational chromatin-state framework as follows:

- Active promoters: H3K4me3 ± H3K27ac within 1 kb of a TSS
- Bivalent promoters: H3K4me3 + H3K27me3
- Active distal enhancers: H3K4me1 ± H3K27ac outside promoter-proximal regions
- CTCF-associated insulators: CTCF-positive CREs not otherwise classified above
- Unresolved: CREs lacking the above chromatin-state signatures.

All overlaps required at least 1 bp.

### Repeat content and mapping ambiguity

Repeat content was quantified for each CRE using RepeatMasker annotations.^48^ CREs were classified as low (≤25%), moderate (>25% to ≤75%), or high (>75%) repeat content based on the fraction of bases overlapping repetitive sequences. To estimate mapping ambiguity, 10,000 CREs were sampled from each repeat-content class and remapped to the genome; the fraction of CREs with multiple genomic alignments was used as a measure of mapping ambiguity.

### MPRA overlap

To evaluate functional activity of annotated CRE classes, we intersected the cortical CRE map with a published massively parallel reporter assay (MPRA) dataset.^58^ Only a subset of cortical CREs overlapped tested MPRA elements due to the small size of the MPRA dataset, however nearly all MPRA sequences were found in our cortical CREs. Functional outcomes were summarized for these overlapping subsets and classified as activator, silencer, or no effect, according to the external MPRA annotations.

### CRE–gene linkage with C. Origami

To assign candidate target genes to cortical CREs, we used C.Origami,^70^ a multi-modal deep neural network that performs *de novo* prediction of cell-type-specific chromatin organization using DNA sequence and cell-type-specific genomic features. Combined ATAC-seq BAM files and CTCF CUT&Tag BAM files were processed according to the C.Origami workflow (https://github.com/tanjimin/C.Origami/). C.Origami generated gene-centered chromatin contact prediction windows extending across local genomic neighborhoods. From the resulting CRE–gene linkage outputs, we retained three summary tables: (1) CRE-level summaries, including the number of linked genes, nearest gene, and strongest linked gene (per C.Origami chromatin contact predictions); (2) individual CRE–gene links with linkage scores and distances to TSS; and (3) gene-level summaries of the number of linked CREs per gene. CRE–gene linkage architecture was summarized by: (i) whether the strongest linked gene matched the nearest gene, and (ii) the number of genes linked per CRE. These linkage summaries were integrated into the merged cortical CRE dataframe with by genomic coordinate overlap (chromosome, start, end) for downstream comparisons across annotation, donor recurrence, developmental persistence, lineage sharing, and sex bias.

### Donor frequency and developmental persistence

To quantify recurrence (CRE sharing) across individuals, a donor-by-CRE presence/absence matrix was generated from donor-specific pseudo-bulk peak calls. For each CRE, we counted the number of donor-specific maps in which the CRE was detected (donors_present). CREs were then grouped into donor recurrence classes by number: 0, 1, 2–4, 5–9, 10–19, 20–39, 40–53, and 54 donors. To quantify developmental persistence, we similarly generated gestational-week-resolved presence/absence profiles across GW16–GW24 and counted the number of gestational weeks in which each CRE was detected (weeks_present). CREs were grouped by number as 0, 1, 2–4, 5–8, or 9 weeks. Lineage sharing was defined analogously as the number of major cortical lineages in which a CRE was detected (lineages_present) and grouped by number as 1, 2–4, 5–6, or 7 lineages.

### Differential accessibility analysis

Sex differential accessibility analysis was performed using DESeq2.^83^ Read counts for each donor-by-cell-type pseudo-bulk were generated using featureCounts from the Subread package.^84^ We then fitted a generalized linear model using donor sex, donor age, and cell-type as covariates. CREs were classified as male-biased, female-biased, or unbiased based on adjusted p-value thresholds and the sign of the log_2_ fold-change. In addition to differential accessibility, threshold-based overlap categories were defined as male only, female only, or both sexes according to whether a CRE was present in sex-stratified pseudo-bulk maps.

### TF sequence grammar and candidate regulator prioritization

To characterize transcription factor (TF) sequence grammars, we applied a deltaSVM-based^46^, a machine learning-based motif discovery framework to the merged cortical and lineage-specific CRE maps. Index-script outputs were converted to narrowPeak format, and sequence quality and training sets were generated using gkmQC^43^ with default parameters (10-mer word length; 5,000 CREs per training group). Positive and negative training sequences from the top 100,000 peaks were then used to train LS-GKM models with word length 11.^6^ Trained models were used to score all non-redundant 11-mers generated using the gkm-pipeline tool (https://github.com/Dongwon-Lee/gkm-pipeline/). Scores were generated for the merged cortical map and for each lineage-specific map, and were z-score normalized within each set. Cis-BP 2.0 human TF motifs were downloaded from the MEME Suite.^85^ Motifs were matched to 11-mers using FIMO-based scanning of 11-bp windows across each motif. TF motif enrichment was quantified as previously described,^45^ as the relative enrichment of motif-matching k-mers among the top 5% of scored 11-mers. For lineage-specific candidate regulator prioritization, TF motif support was integrated with lineage-matched TF gene expression. Within each lineage, motif scores and expression values were z-score normalized and summed to generate a combined support score. Lineage selectivity was then quantified as the difference between the highest lineage-specific combined score and the mean combined score across the remaining lineages. Representative lineage-selective TF candidates were prioritized from this integrated score.

### Statistical summaries and visualization

All summary statistics and visualizations were generated in R using ggplot2, dplyr, data.table, and related packages. Heatmaps, line plots, ridge plots, violin plots, and summary dot plots were used to visualize recurrence, persistence, annotation composition, motif candidates, and sex-biased CRE properties. Unless otherwise stated, percentages shown in compositional figures were normalized within the class indicated on the x-axis or the legend.

## SUPPLEMENTARY FIGURE LEGENDS

**Figure S1.**
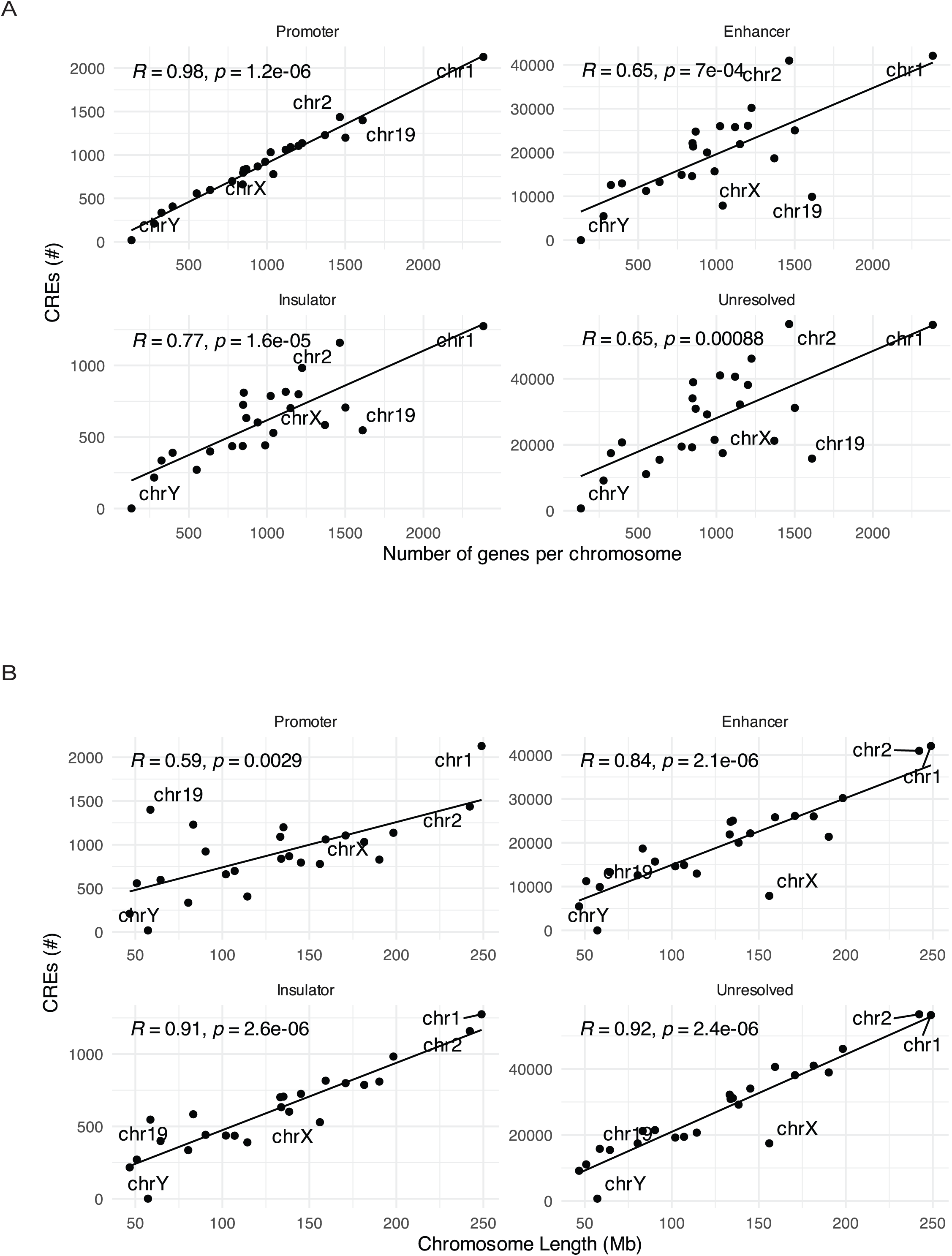
Chromosome-level abundance of cortical CRE classes relative to gene numbers. **(A)** Chromosome-level counts of cortical CREs compared with chromosome gene number for major annotation classes. Promoter-associated CREs showed the strongest relationship to gene abundance, whereas enhancer, insulator, and unresolved CRE classes showed weaker correlations. **(B)** Chromosome-level counts of cortical CREs compared with the size of each chromosome. Promoter-associated CRE counts showed weaker relationship with chromosome size than with number of genes (as expected). Enhancer, insulator and unresolved CREs showed stronger correlations with chromosome length than with number of genes in a given chromosome. Pearson correlation coefficients are shown in each panel. These analyses indicate that promoter-proximal regulatory elements are more tightly coupled to gene-rich genomic regions than other CRE classes.

**Figure S2.**
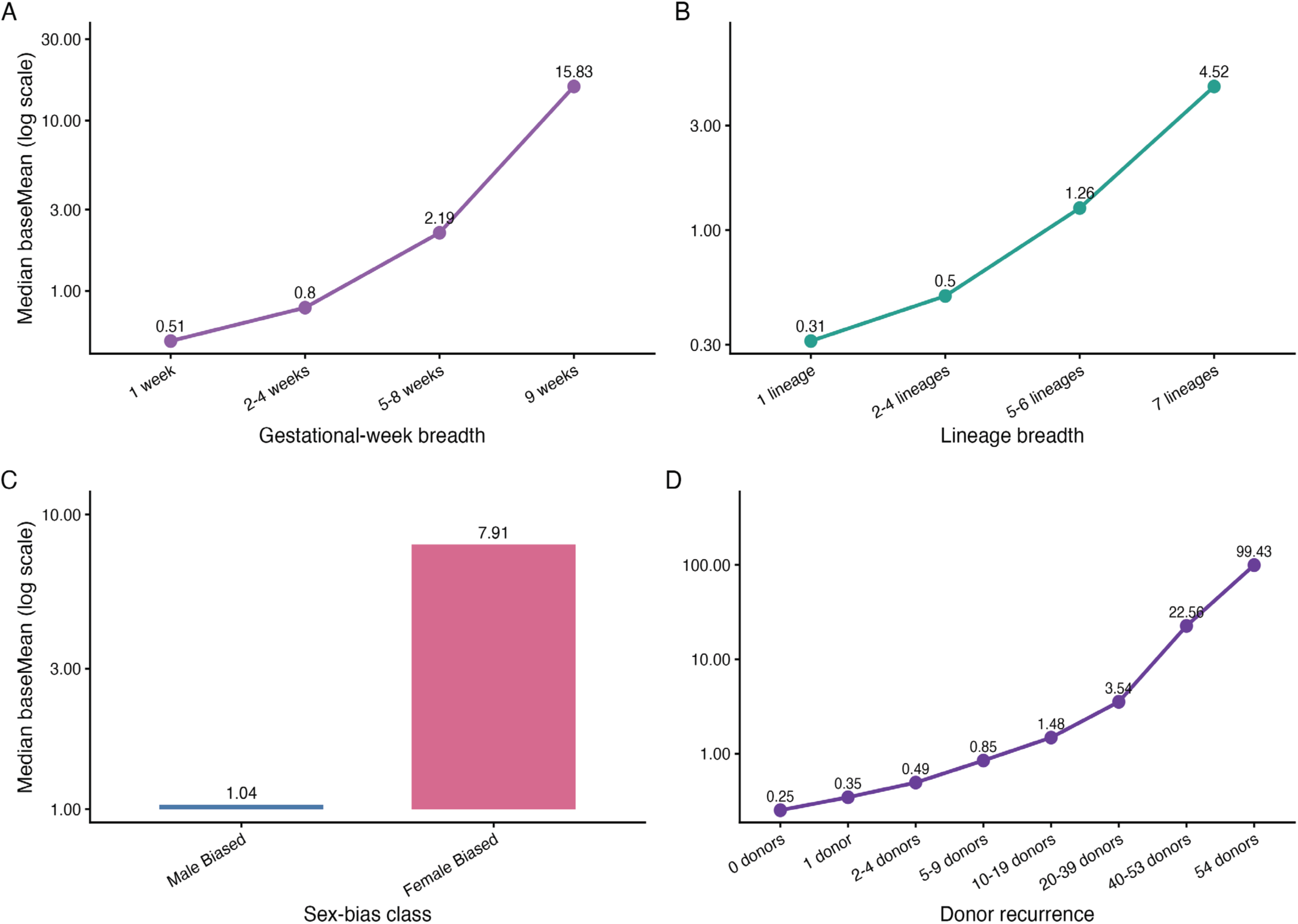
Normalized accessibility activity tracks developmental persistence, lineage sharing, sex bias, and frequency. Median normalized accessibility (baseMean) was calculated across CRE classes defined by gestational-week sharing, lineage sharing, sex bias, and donor recurrence. **(A)** CREs detected across increasing numbers of gestational weeks showed progressively higher median baseMean, consistent with greater signal strength among developmentally persistent elements. **(B)** CREs shared across increasing numbers of cortical lineages likewise showed higher normalized accessibility than lineage-restricted CREs. **(C)** Female-biased CREs had substantially higher median normalized accessibility than male-biased CREs. **(D)** Donor recurrence was strongly associated with normalized accessibility, with broadly recurrent CREs showing markedly higher median baseMean than donor-restricted CREs. Together, these panels indicate that the most robust CREs form a recurrent, broadly shared regulatory core, whereas restricted CREs are weaker on average.

## Notes

### Competing Interest Statement

The authors have declared no competing interest.

### Funding

- Simons Foundation

